# BrainVAE: Exploring the role of white matter BOLD in preclinical Alzheimer’s disease classification

**DOI:** 10.1101/2025.05.07.652697

**Authors:** Yikang Li, Lyuan Xu, Lianrui Zuo, Yukie Chang, Zhaohua Ding, Adam W. Anderson, Kurt G. Schilling, John C. Gore, Yurui Gao, the Alzheimer’s Disease Neuroimaging Initiative

## Abstract

**INTRODUCTION:** Like gray matter (GM), white matter (WM) BOLD functional signals change in preclinical AD. However, the potential of WM BOLD for identifying preclinical AD remains underexplored.

**METHODS:** We developed BrainVAE, a transformer-based variational autoencoder with interpretability, to classify preclinical AD and normal controls using resting-state fMRI data. We benchmarked BrainVAE against nine alternative models under three input configurations: WM-only, GM-only, and combined WM+GM. Interpretability analysis was also performed to investigate each brain region’s contribution to the classification task.

**RESULTS:** BrainVAE outperformed other models and performed well (accuracy = 83.42%, F1-score = 91.62%, AUC = 64.50%) using the combined input compared to WM-only and GM-only. Specific WM bundles--including corpus callosum, fornix, and corticospinal tract—were among the most influential features contributing to the classification.

**DISCUSSION:** Incorporating WM BOLD signals improves the distinction of preclinical AD from controls, underscoring the potential role of WM BOLD features for detecting early-stage AD.

**Highlights:**

- BrainVAE integrates white and gray matter BOLD signals for classification of pre-AD and controls.
- BrainVAE achieves high accuracy (83.42%) and F1-score (91.62%) in identifying pre-AD.
- Models using combined WM+GM inputs outperform those using WM-only or GM-only inputs.
- WM regions, such as corpus callosum and fornix, contribute significantly to model predictions.
- Results suggest WM BOLD signals are informative markers for early AD detection.

## 1. Background

Alzheimer’s disease (AD) is a progressive neurodegenerative disorder characterized by the accumulation of amyloid-beta (Aβ) plaques and hyperphosphorylated tau proteins, leading to widespread neuronal dysfunction and ultimately cognitive decline^1^. Functional alterations may begin years or even decades before the onset of clinical symptoms, involving both gray matter (GM) and white matter (WM) networks.^2^ This prolonged preclinical phase offers a critical window for early detection of AD.

Resting state functional MRI (fMRI) has emerged as a promising tool for detecting early functional disturbances in the brain.^2,3^ However, most prior fMRI research has focused on GM signals, often discarding signals in WM as nuisance variables.^4,5^ Recent evidence suggests that WM BOLD fluctuations—although weaker—are reliably detectable and associated with brain activity. Importantly, our work has demonstrated that fMRI measures in WM are altered not only in mild cognitive impairment and AD dementia^6^ but also during the preclinical AD period^7^, suggesting their potential for early disease classification.

Despite these promising observations, it remains unclear whether WM fMRI signals provide additional diagnostic power beyond traditional GM fMRI features alone. We hypothesized that adding WM functional information to GM would enhance performance in identifying preclinical AD. However, testing this hypothesis using traditional methods poses several critical challenges. First, WM BOLD signals inherently possess lower signal-to-noise ratios than GM^8,9^, while the functional parcellation of WM remains less mature compared to GM, making it challenging to compute robust and reproducible metrics of functional connectivity. Second, functional connectivity matrices in preclinical AD are characterized by subtle, small-magnitude changes that are distributed across many ROI pairs, requiring models capable of detecting weak and spatially diffuse inter-regional dependencies.^10,11^ Third, the relatively small number of preclinical AD datasets leads to class imbalance, increasing the risk of overfitting, particularly for standard supervised learning approaches. In addition, recent developments in deep learning provide interpretable attention mechanisms that enable the quantitative assessment of each white matter bundle’s contribution to classification outcomes^12^.

To address these challenges, we propose **Brain Variational AutoEncoder (BrainVAE)**, a novel deep learning model designed to integrate WM and GM fMRI measures for the classification of preclinical AD and controls. First, we employ deep neural networks capable of extracting robust and biologically relevant features from fMRI data with low signal-to-noise ratios and anatomical variability, particularly within WM. Second, we use a transformer-based encoder architecture, whose self-attention mechanism effectively captures subtle and spatially diffuse inter-regional dependencies between brain regions.^13,14,15^ This approach enhances the model’s generalizability while maintaining biological interpretability, enabling accurate identification of preclinical AD based on integrated WM and GM functional connectivities.

BrainVAE extends beyond conventional classification models by integrating interpretability directly into its architecture. Through the transformer-based encoder, BrainVAE systematically assigns attention weights to individual GM parcels and WM bundles, enabling identification of brain regions, which is most critical for distinguishing preclinical AD from normal aging. To validate the effectiveness of our approach, we benchmarked BrainVAE against a diverse range of machine learning and deep learning models. Each model was evaluated using three types of input datasets: GM-only, WM-only, and combined GM-WM functional connectivity matrices. This experimental design allowed us to assess both the overall classification performance and the added diagnostic value of incorporating WM signals. Overall, BrainVAE demonstrated robust generalization across input types while providing insights into the functional relevance of WM networks in early AD diagnosis.

## 2. Methods

### 2.1 Data and Participants

The dataset for primary classification was sourced from the Alzheimer’s Disease Neuroimaging Initiative (ADNI), comprising resting state fMRI and T1-weighted images of 78 participants with preclinical AD (pre-AD) (64% female; 77.0 ± 6.8 years) and 217 normal controls (NC) (56% female; 75.2 ± 7.4 years). All these participants were clinically unimpaired, defined by a Clinical Dementia Rating (CDR) of 0 and a Mini-Mental State Examination (MMSE) score between 24 and 30. Pre-AD participants were Aβ-positive, meeting at least one of the following criteria: cerebrospinal fluid (CSF) Aβ < 459 pg/mL^16^, AV-45 PET standardized uptake value ratio (SUVR) > 1.22^17^, Pittsburgh compound B (PiB) PET SUVR > 1.42^18^, or Florbetaben (FBB) PET SUVR > 1.478^19^. In contrast, NC participants were Aβ-negative across all available modalities.

For pretraining purposes, fMRI and T1-weighted data from 1953 cognitively normal older participants (55% female; 70.8 ± 7.2 years) were obtained from the Baltimore Longitudinal Study of Aging (BLSA)^20^ and the Open Access Series of Imaging Studies (OASIS)^21^. These datasets have comparable spatial resolution, repetition time and number of time points.

Detailed cohort characteristics, including demographic information and clinical measures, are summarized in Table 1.

**Table 1.**
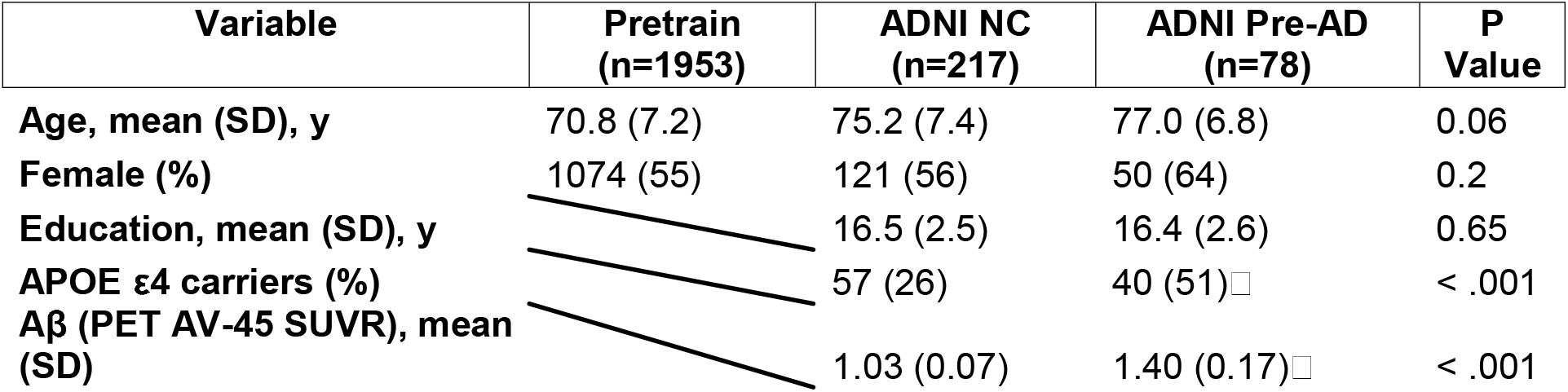
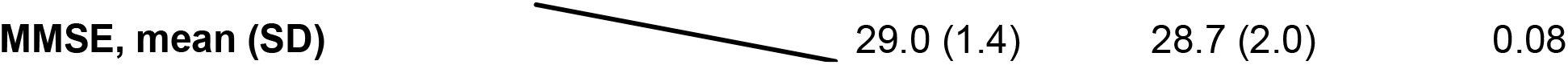
Characteristics of the Study Cohort including classification datasets and pretrain datasets. Note: ^a^ Uncorrected *P* < .05 ^b^ FDR-corrected *P* < .05.

### 2.2 Data Processing

Data processing includes image preprocessing and evaluation of FC matrices, briefly stated as below (see ref ^7^ for more details).

The fMRI images were first corrected for slice timing and head motion; participants with head rotation > 2° or head translation > 2mm were excluded. Twenty-four motion parameters and the mean signal from CSF were regressed out^22^. Each time series was detrended and bandpass filtered (0.01– 0.1□Hz). These preprocessing steps were executed using the Data Processing Assistant for Resting-State fMRI toolbox^23^. Meanwhile, tissue probability maps (TPMs) for brain-wide GM, WM, and CSF were generated by segmenting the T1-weighted images using the Computational Anatomy Toolbox (CAT12)^24^. Both the filtered fMRI data and TPMs were normalized to MNI space. To prevent partial volume effects at the WM-GM interface, spatial smoothing was intentionally omitted.

GM and deep WM were parcellated into 200 cortical parcels and 48 WM bundles, based on the Schaefer atlas^25^ and the JHU’s Eve atlas^26^, respectively, resulting in a total of 248 regions (Fig. 1a). Regional time series were extracted by averaging the z-scored time series within each region. FC was computed using Pearson’s correlation between the time series of each pair of regions, yielding a 248□×□248 FC matrix for each participant. To curb the spurious correlations arising from noise alone, FC values below 0.1 were set to zero.

**Figure 1.**
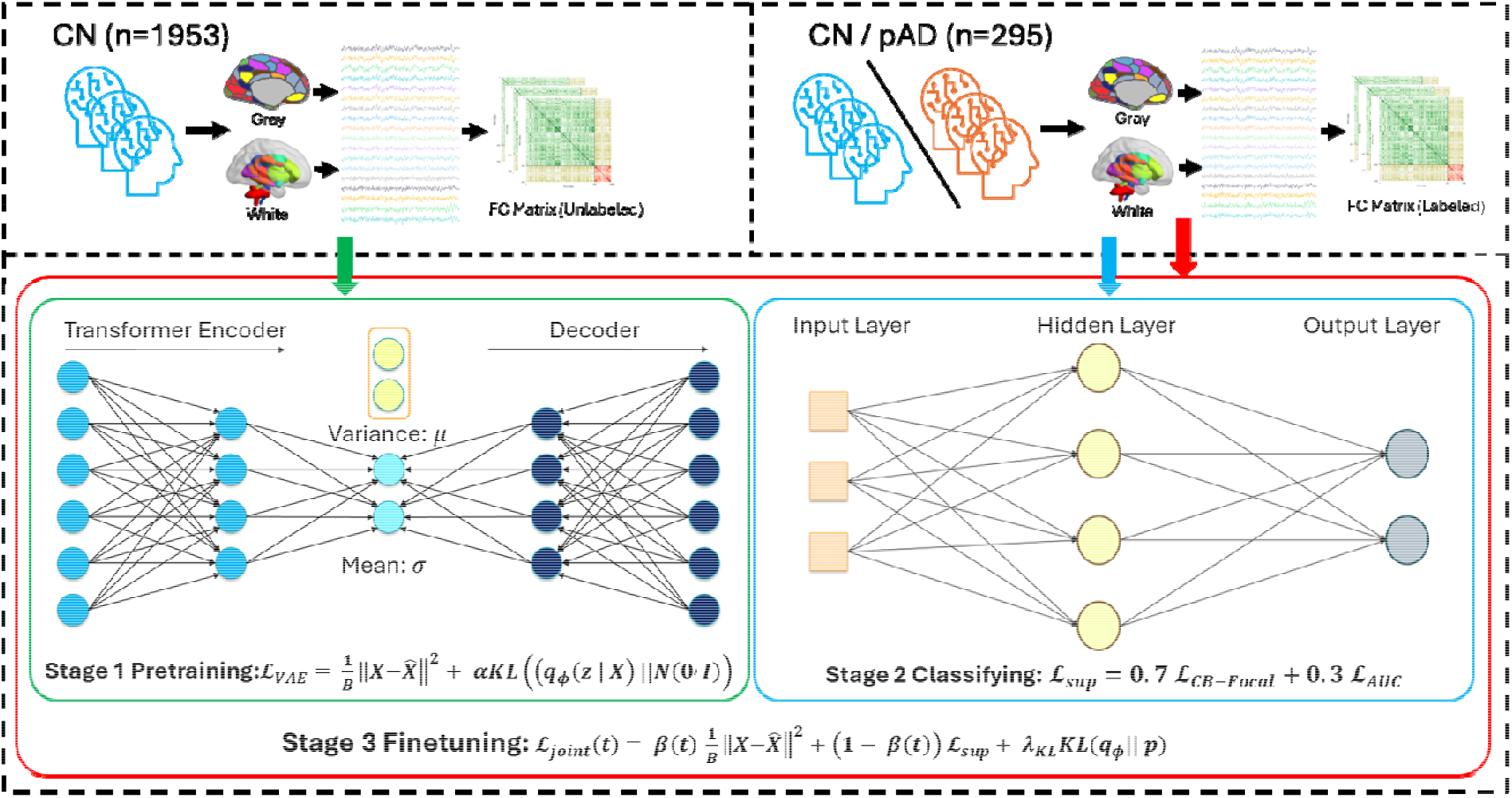
The overall pipeline of BrainVAE consists of three stages: pretraining, supervised-learning and joint finetuning. The first datasets on the upper left are the large healthy datasets that are used for pretraining stage. The labeled datasets were used in the supervised-learning and joint finetuning stages.

**Figure 2.**
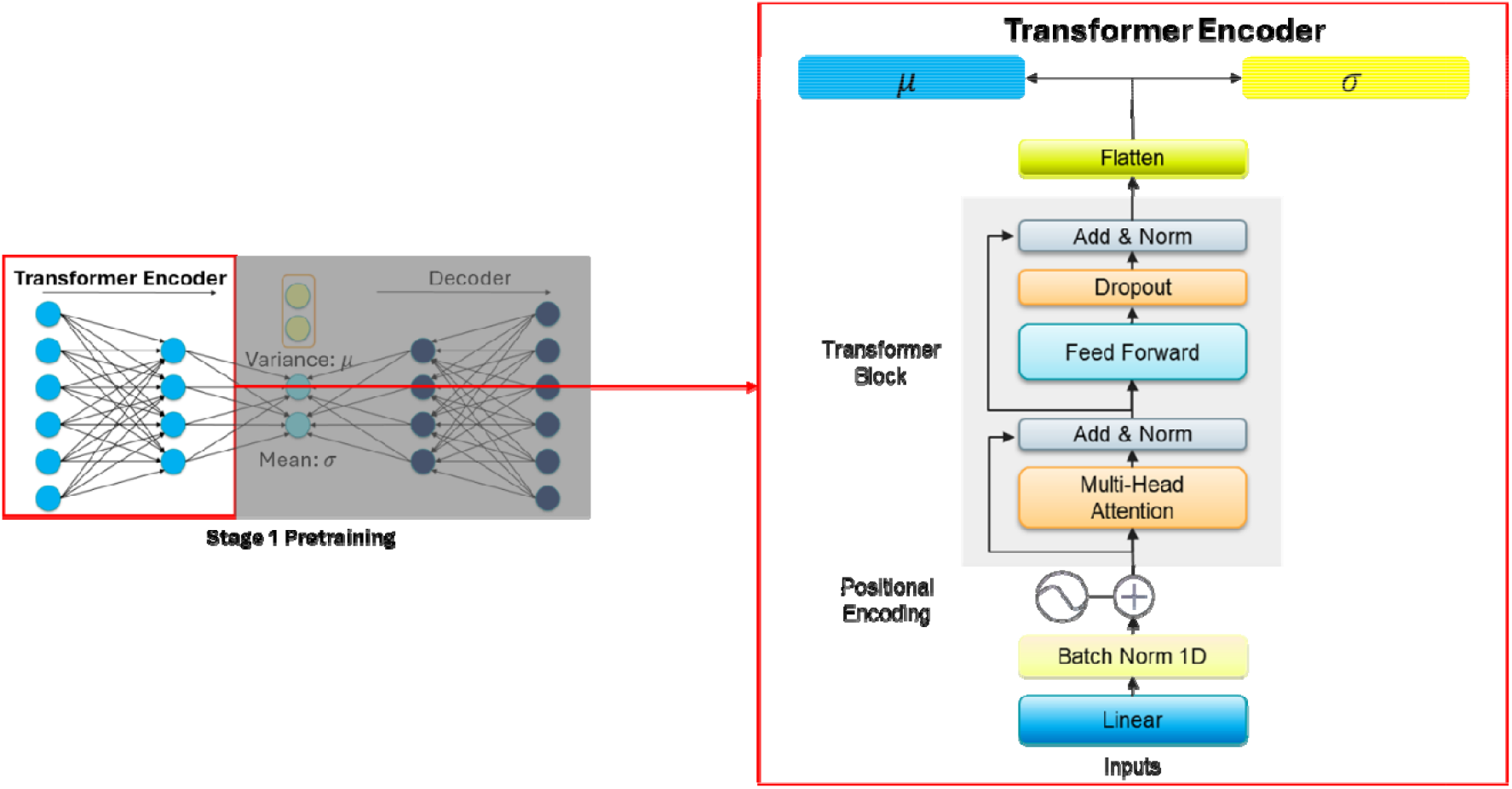
The main architecture of the Transformer-based encoder in the BrainVAE model.

### 2.3 BrainVAE Model

FC matrices are derived from datasets with significant class imbalance, as the number of preclinical AD scans is substantially lower relative to cognitively normal controls. Under these circumstances a discriminative neural network is prone to over□fitting the minority class and to ignoring subtle but physiologically meaningful signals between-group difference. We therefore adopt a variational auto□encoder framework, which first learns—without labels—a low□dimensional, smooth manifold representing normal GM and WM connectivity. Supervised classification is performed only in that compressed latent space; these yields strong regularization, an explicit probabilistic measure of abnormality via the latent Kullback–Leibler (KL) term, and a principled way to combine unsupervised information from 1,953 healthy controls with the limited labelled cohort.

The neural network, shown in Figure 1, comprises a transformer□based encoder that converts a 248□×□248 correlation matrix,□*X*, into latent mean *µ* and log-variance log (*σ* ^2^), a decoder that reconstructs *x* from a stochastic latent vector *z*, and a multilayer perceptron classifier that maps *s* ∈ ℝ ^2^.

To stabilize learning under class imbalance (78□pre-AD vs 217□CN), network parameters are optimized in three successive stages described in Section 2.3.3.

#### 2.3.1 Pretrained Variational AutoEncoder

In this study, the input to the VAE is the correlation matrix derived from each participant’s resting-state fMRI signals. For each participant we observe a symmetric correlation matrix:

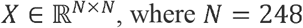

The goal is to encode the matrix into a latent vector *z* ∈ ℝ^d^, *d* = 64, distributed according to the Gaussian prior *p*(*z*) = N(0, I), and then decode *z* back into a reconstructed correlation matrix 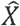 with the least difference compared to *X*. During the pretraining, the network is optimized so that the reconstructed output is close to the original input, but with additional constraints that ensure the latent space remains well-organized and continuous.

The encoder portion of the VAE applies several learnable transformations to the correlation matrix. The encoder parameterized by *ϕ* approximates the posterior distribution:

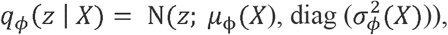

where *µ*_ϕ_ and log 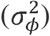 are learned functions mapping the input FC matrix *X* ∈ ℝ^*N*×*N*^ to a d-dimensional latent space, representing the mean and variance of the Gaussian distribution, respectively. Instead of directly sampling *z* from these learned distributions (which would break the computational graph), a reparameterization step is introduced. In this step, the network draws random noise *ϵ* from a standard normal distribution and constructs

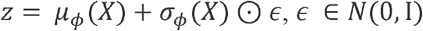

so that the backpropagation algorithm can still update all parameters.

The decoder *p*_ψ_ (*X*|*z*) receives *z* and attempts to reconstruct the correlation matrix

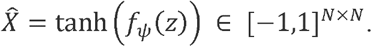

Since the reconstruction target is correlation coefficient, which has values between -1 to 1, we used a hyperbolic tangent as the final activation layer to enforce this data range. Reconstruction fidelity is measured with a mean-squared error (MSE) or similar criterion, ensuring that reconstructed matrices approximate the actual functional connectivity patterns in the training dataset. To keep the latent variables from collapsing onto arbitrary points, an additional penalty term known as the Kullback– Leibler (KL) divergence is included. The KL term forces the learned distribution *N*(*µ, σ*^2^) to remain close to a chosen prior (typically *N*(0,*I*)). Thus, the overall training objective for stage 1 can be written as:

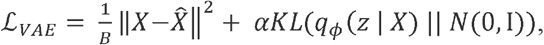

where B is the mini batch size, and *α*= 1. The first term encourages accurate reconstruction; the KL term constrains the latent distribution to remain close to the unit Gaussian, preventing “holes” where the decoder has never been trained and thus enforcing a smooth, informative latent manifold.

#### 2.3.2 Transformer-based Autoencoder

Traditional multilayer perceptron (MLP) encoders treat each input as an independent feature and rely on fully connected layers to capture relationships across regions, which can be inefficient or ineffective for modeling complex inter-regional dependencies in brain connectivity data. To better capture the structured relationships inherent in functional connectivity (FC) matrices, we adopted a transformer-based encoder architecture. Transformers are suited for this task due to their self-attention mechanism, which explicitly models pairwise interactions between all regions of interest (ROIs), regardless of spatial proximity or ordering. By treating each row of the FC matrix (i.e., each ROI’s connectivity profile) as a token, the transformer can learn contextualized representations that reflect both local and global connectivity patterns. This structure-aware encoding is especially valuable in the context of preclinical AD, where subtle but distributed alterations in connectivity may serve as early biomarkers. The transformer encoder was used for both the unsupervised pretraining phase and the downstream classification task.

The inputs for the transformer-based encoder are the rows of *X* interpreted as 248 ordered tokens {*x*_1_,*x*_2_, …, *x*_*N*_}∈ ℝ^*N*^. Each token is linearly embedded 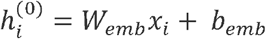, where *W*_*emb*_ ∈ ℝ^*d* × *N*^, *d* is the dimensionality of each query or key, followed by batch normalization across tokens. Then two identical self-attention blocks (dropout□0.3) are stacked. For a set of token features *H* ∈ ℝ^*d* × *N*^, each attention head *h* computes 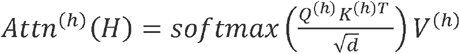, with learnable projections 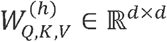 where queries: 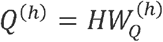, keys: 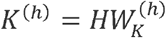 and values: 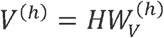.

Our transformer based encoder has four heads, which run in parallel; their outputs are concatenated, passed through a 256-unit feed-forward network, and combined with residual connections and layer-norm—this constitutes one transformer block, yielding a depth sufficient to model higher-order connectivity patterns while avoiding over-fitting.

Our transformer-based encoder enables the latent representation *z* to encode not only the overall covariance structure but also specific region-to-region dependencies that may be highly relevant for distinguishing preclinical AD from normal aging. In the final classification step, the network extracts z and feeds it into a supervised head (a series of fully connected layers) trained to categorize each participant as either cognitively normal or pre-AD.

#### 2.3.3 Three □ stage training protocol

The main architecture of BrainVAE leverages the strengths of both unsupervised representation learning and attention-driven encoding. In this case, to maximize the strength of representation learning and avoid the classic label imbalance problems, we stated the three stages training protocol within the BrainVAE model. The first stage is unsupervised pre-training the transformer-based encoder and decoder. During the process, parameters (*ϕ, ϕ*) are learned by minimizing ℒ _*VAE*_ on 2□000 healthy controls. Weights yielding the lowest validation loss are retained.

In the second stage, the classifier is trained on the labeled classification datasets while the encoder and decoder are frozen. The supervised loss is defined as the combination of a class-balanced Focal term^26^ ℒ _*CB*−*Focal*_ and the differentiable AUROC surrogate^27^ ℒ _*AUC*_ :

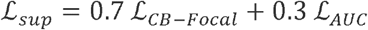

In the third stage, all parameters of encoder, decoder and classifier are trained in a joint loss function for fine tuning. The composite loss is defined as:

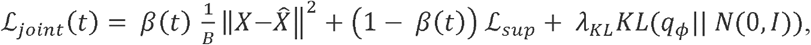

where *λ* _*KL*_ =10^−3^ and 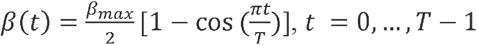, *β*_*max*_ =0.3, and *T* is the number of epochs.

### 2.4 Benchmark Models and Performance Assessments

To validate the effectiveness of the proposed model, a comprehensive set of benchmark models was implemented for comparisons, including classic machine learning algorithms, convolution neural networks (CNN)-based models, and graph neural networks (GNN)-based models (Fig. 3). Specifically, the machine learning models included support vector machines (SVM)^30^, multilayer perceptron (MLP)^31^, and random forests^32^, all of which used regional time series as input. The CNN-based models included BrainNetCNN^33^, Transformer^34^ and Brain Network Transformer^14^, using FC matrices as input. The GNN-based models included BrainGNN^35^, interpretable GNN (IBGNN)^36^ and its variant IBGNN+. These models operate on graph representations, where nodes represent brain regions and edge weights encode inter-regional functional connectivity.

**Figure 3.**
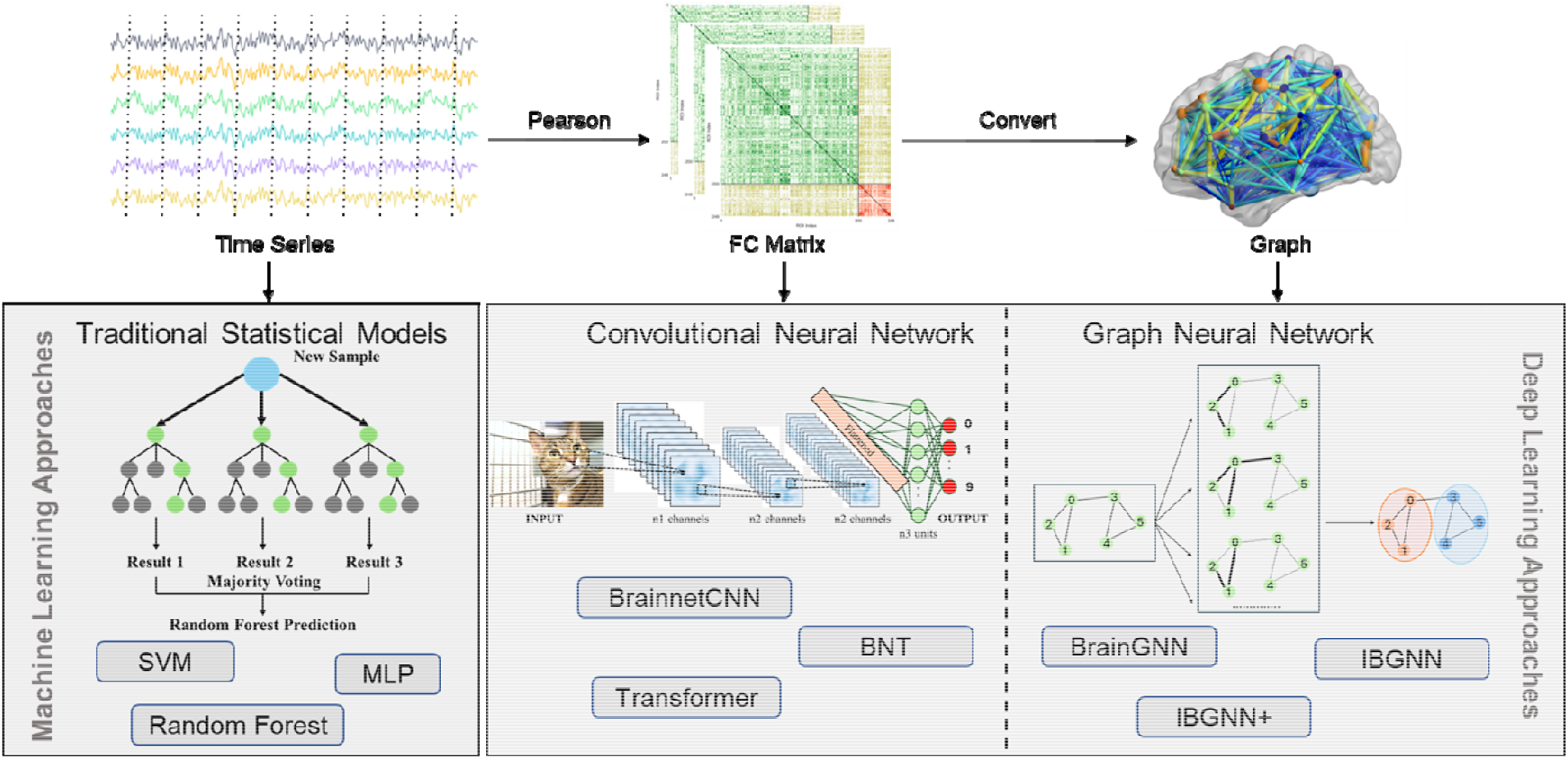
The benchmarked models across different types of data and different approaches.

Model performance was assessed using five-fold cross-validation. with accuracy, F1-score, and receiver operating characteristic curve (AUC) as assessment metrics. Each model was evaluated on three input configurations: one with only GM parcels, one with only WM bundles, and one combining both. This experimental design aimed to assess whether adding WM features improved classification performance.

### 2.5 Interpretability Analysis

The BrainVAE framework incorporates a transformer-based attention encoder, which not only boosts classification accuracy but also provides interpretability by highlighting the most informative brain regions across the cohort. Specifically, the attention mechanism learned to assign higher attention weights to brain regions—either GM parcels or WM bundles—that contribute more to the classification of preclinical AD and NC. The regional importance scores were extracted by averaging the attention weights across all self-attention heads. Higher importance score indicated a larger contribution to the classification.

In the combined configuration (WM+GM), attention scores were specifically examined to quantify how integral WM bundles were to the network’s understanding of emerging pathology. The presence of WM bundles among the top-ranked features provided evidence supporting the critical role of WM signals in early disease detection.

## 3. Results

### 3.1 BrainVAE benchmarking

BrainVAE achieved averaged classification accuracies of 0.7763, 0.8271, and 0.8342 for the WM-only, GM-only and combined WM+GM configurations, respectively, outperforming all nine benchmark models (Table 2). It also yielded the highest average F1-scores for all three configurations: 0.873 (WM-only), 0.904 (GM-only) and 0.916 (WM+GM). In terms of AUC, BrainVAE ranked first for the WM-only and WM+GM configurations, and among the top three for GM-only. Collectively, these results suggest that BrainVAE model provides superior overall classification performance (Table 2).

**Table 2.**
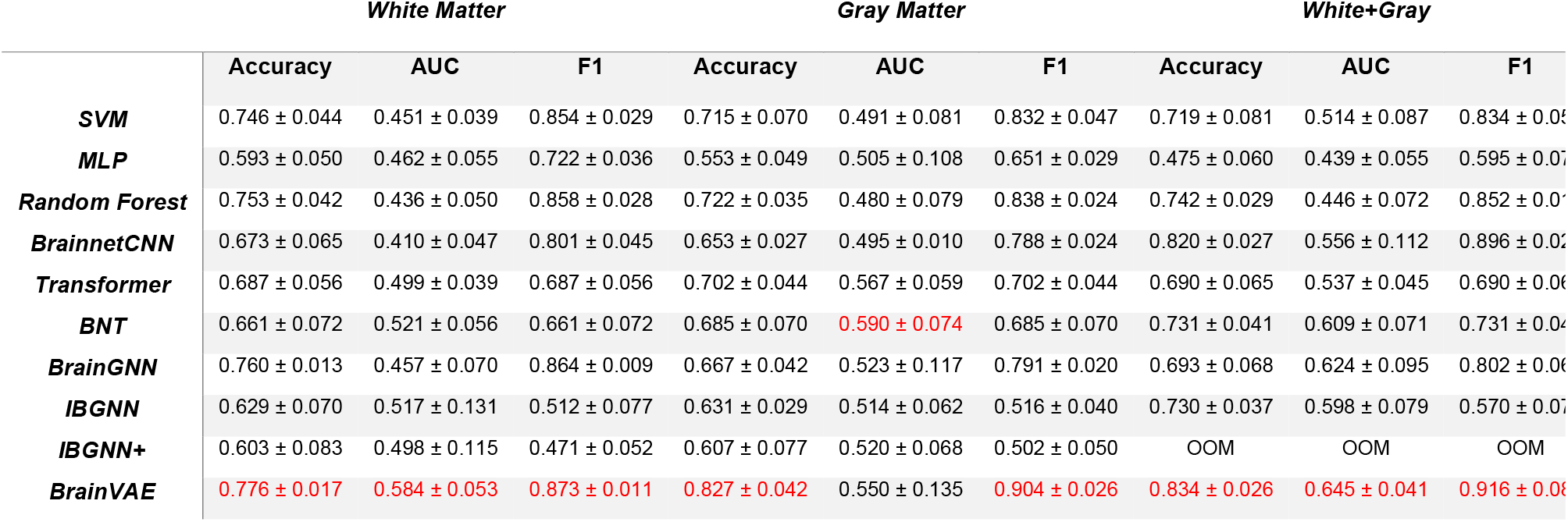
The performance comparison between BrainVAE and other models with different types of data input.

Using only WM BOLD features, BrainVAE achieved a 6% higher AUC, 6% lower accuracy and 3% lower F1-score compared to the GM-only inputs, suggesting that WM BOLD carries at least comparable information to conventional GM BOLD for classification. Notably, all benchmark models showed lower or almost the same AUCs when using WM-only versus GM-only input (Table 2), indicating BrainVAE may be uniquely more sensitive to WM BOLD information.

### 3.2 Classification with combined WM and GM features

Across nearly all models, performance was improved when using combined WM+GM features as input, compared to using either WM-only or GM-only input alone. The combined input consistently produced the highest accuracy, F1-score, and AUC for most models. Full comparisons across models and input types are presented in Table 2 (and illustrated in Fig. 4).

**Figure 4.**
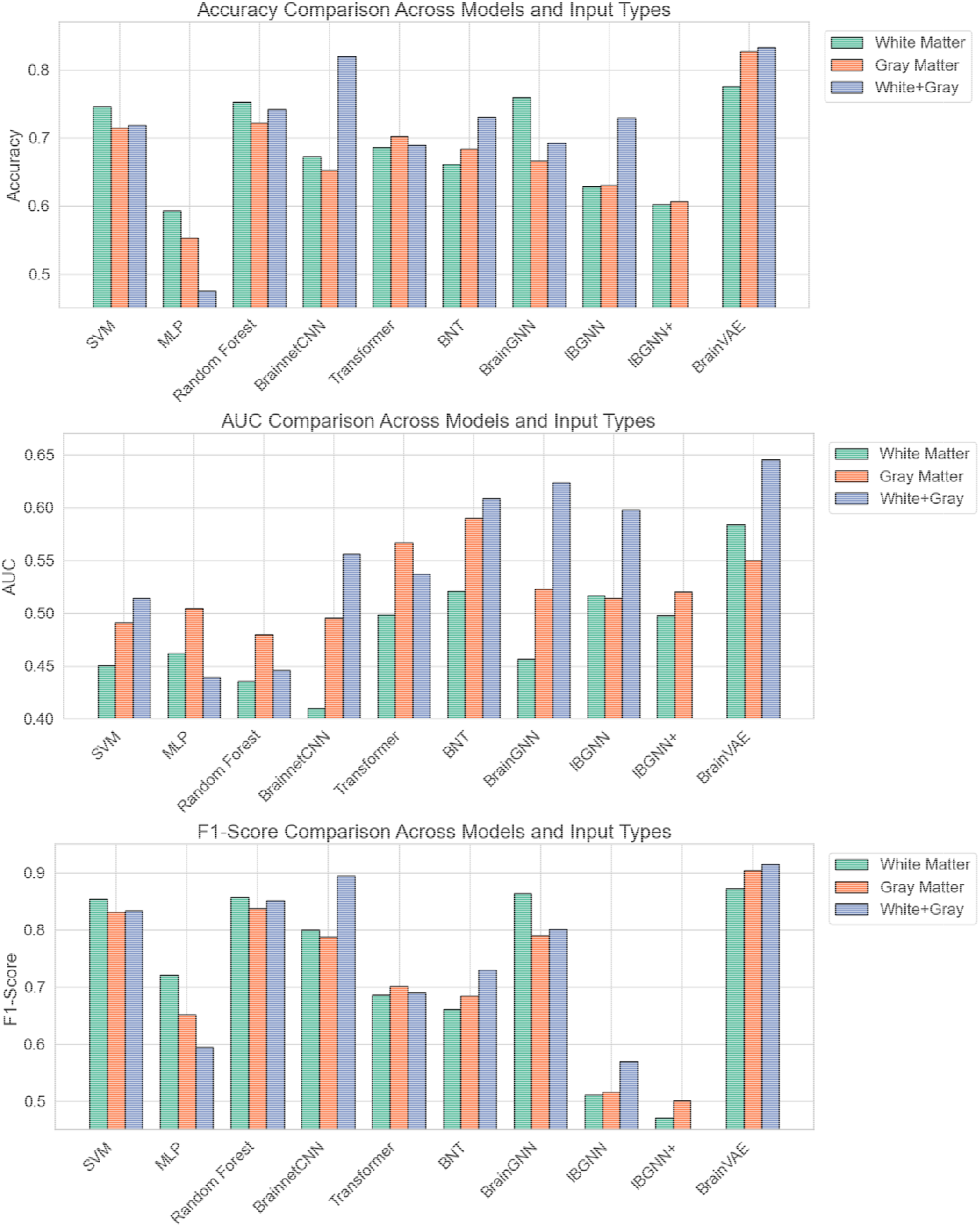
Visualization of classification performance (Accuracy, AUC, and F1-score) for BrainVAE and baseline models across three input conditions: white matter only, gray matter only, and combined matter.

### 3.3 Interpretability Analysis Results

Figure 5A presents the resulting attention score matrix for the combined WM+GM input, where rows and columns correspond to 248 ROIs (200 GM, 48 WM).

**Figure 5.**
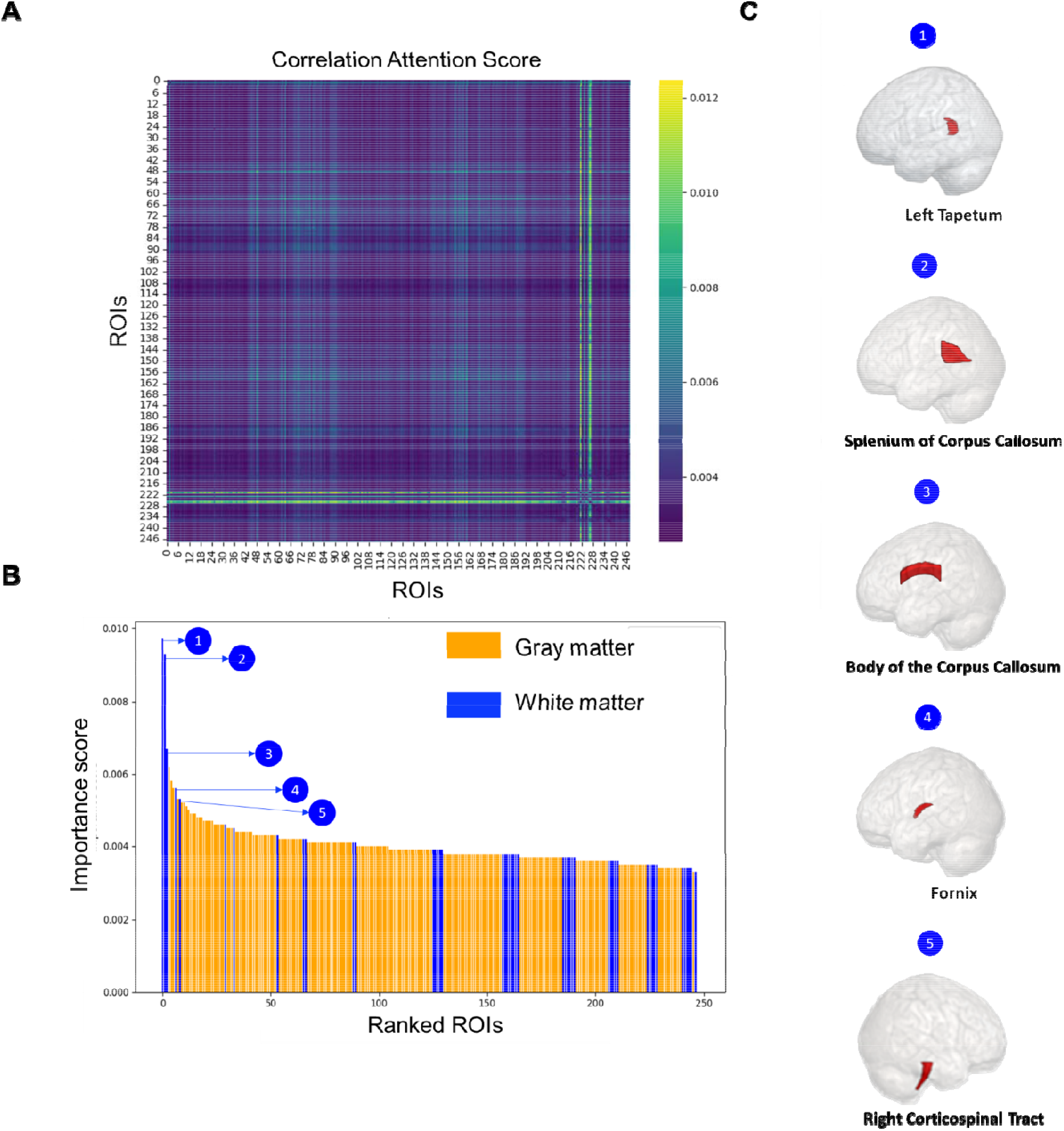
A. Correlation attention scores across combined gray and white matter ROIs, indicating network-level contributions to classification. B. Ranked importance scores for each ROI, with gray matter (orange) and white matter (blue), highlighting the top five white matter ROIs. C. Top five important white matter ROIs identified by BrainVAE’s attention mechanism for pre-clinical Alzheimer’s disease classification.

To evaluate the importance of each ROI, we summed the attention weights across each row in the matrix. The resulting importance distribution is shown in Figure 5B, with GM and WM regions color-coded in orange and blue, respectively. Several WM bundles ranked among the highest in attention score, with some exceeding all GM parcels. The top-ranked WM regions were further visualized in Figure 5C, including left tapetum, splenium of corpus callosum, body of corpus callosum, fornix, and right corticospinal tract.

## 4. Discussion

The main goal of this study is to investigate whether incorporating WM functional signals, alongside the more commonly studied GM connectivity patterns, could enhance the classification of individuals with pre-AD versus normal controls. Leveraging a novel transformer-based variational autoencoder, BrainVAE, we systematically evaluated classification performance across multiple input configurations and model types. Three key findings emerged: (1) BrainVAE consistently outperformed all benchmark models in distinguishing pre-AD from cognitively normal controls; (2) combining WM+GM input led to improved classification accuracy across a range of models; and (3) attention-based interpretability revealed that several WM bundles were among the most influential regions contributing to model predictions.

### 4.1 Overall Performance of BrainVAE

BrainVAE achieved the highest performance in distinguishing preclinical AD from normal controls, demonstrating its ability to extract informative functional features for classification even if no between-group differences were observed in clinical cognitive scores (Table 1).

Classification accuracy remained consistent across different input conditions, including WM-only (0.7763), GM-only (0.8271), and combined WM+GM (0.8342), with relatively small performance gap. This suggests that BrainVAE is capable of effectively leveraging information from WM connectivity, even in face of its lower signal-to-noise ratio and less mature functional parcellation. In addition, the model produced stable results across cross-validation folds, as reflected by the small standard deviations reported in Table 2, indicating robustness to class imbalance and limited sample sizes in the pre-AD group.

Across all benchmark models, BrainVAE consistently achieved the highest classification performance under each input condition, including WM-only, GM-only, and combined GM-WM connectivity. This suggests that its transformer-based encoder and variational autoencoder structure provided stronger representational capacity for capturing informative features in the functional connectivity data.

### 4.2 Value of WM BOLD signals

Not only BrainVAE, but also nearly all benchmark models demonstrated improved classification performance when using combined WM+GM input compared to GM-only or WM-only, regardless of model architecture. While GM-only inputs generally outperformed WM-only, the combined input consistently yielded the highest accuracy, F1-score, and AUC across all model categories. These findings highlight the added diagnostic value of incorporating WM BOLD features and suggest that its contribution is robust across both classic machine learning and deep learning frameworks.

### 4.3 Interpretability Analysis

The interpretability results revealed that BrainVAE does not distribute predictive attention uniformly across all brain regions, but instead concentrates it on a selective subset of regions, spanning both WM bundles and GM parcels. The distribution of attention weights exhibited a non-random, asymmetric pattern, where a small number of regions—particularly within WM—received disproportionately high scores. This indicates that the model identifies localized functional features as critical for classification, rather than treating all regions as equally informative or relying on uniform global patterns.

The consistent presence of WM bundles among the top-ranked attention regions suggests that the model considers functional interactions within WM as directly informative, rather than as secondary or supporting input. At the same time, known GM regions associated with early-stage AD also received relatively high attention scores, reflecting BrainVAE’s integration of complementary contributions from both tissue types for decision-making.

Together, these findings demonstrate that BrainVAE’s architecture provides a selective focus on functionally meaningful connections and support a regionally specific decision-making strategy that spans both WM and GM domains.

### 4.4 Biological Relevance

The WM bundles identified as critical to BrainVAE’s classification decisions—including the left tapetum, splenium of the corpus callosum, and fornix—are anatomically positioned to support long-range connectivity between cortical regions. The fornix and tapetum are involved in limbic and memory-related processing, while the corpus callosum facilitates interhemispheric information transfer. These tracts contribute to large-scale networks that subserve episodic memory, attention, and executive function—domains known to be subtly affected in the preclinical stage of AD^37^.

The elevated attention weights assigned to these tracts align with accumulating evidence that WM abnormalities may precede or co-occur with cortical atrophy in AD. Myelin loss and axonal disruption in these regions have been linked to Aβ accumulation, particularly during the asymptomatic stage^38,39^. Moreover, alterations in the posterior callosal regions and corticospinal tract have been associated with early functional impairment in motor-related systems, which may reflect broader network disintegration that extends beyond cognitive circuits^40^.

These findings support a growing body of work indicating that functional connectivity in WM is not merely epiphenomenal but may provide early biomarkers for AD detection. Functional MRI studies have shown that BOLD signals in WM are both detectable and behaviorally meaningful, particularly when analyzed in the context of WM–GM integration^41,42^. The identification of high-attention WM bundles by BrainVAE suggests that these regions may capture subtle disruptions in long-range communication that are difficult to detect with GM features alone.

Taken together, these results reinforce a network-based perspective of AD pathology, in which early dysfunction involves distributed alterations across both WM and GM structures, potentially mediated by molecular and structural degeneration within WM tracts.

## 5. Conclusion

In conclusion, this work demonstrates that accounting for WM BOLD features (time series, FC, and graphs) substantially enhanced the detection of preclinical AD. The synergy between WM and GM features underscores the network-centric character of early AD pathology, offering an expanded view beyond cortical atrophy alone. By elucidating which parcels and bundles particularly contribute to classification, the BrainVAE model provides a framework for future efforts in refining early biomarkers and improving disease characterization.

## Acknowledgements

This work was supported by NIH grants RF1MH123201 (Gore and Landman), R01NS113832 (Gore), R21AG083915 (Gao), K01EB032898 (Schilling), I would like to thank my supervisors, Dr. John Gore, Dr. Kurt Schilling and Dr. Yurui Gao for their guidance and support throughout this project.

